# Longitudinal T cell repertoire analysis reveals dynamic clonal T cell populations in Ulcerative Colitis

**DOI:** 10.1101/2025.01.13.632427

**Authors:** Kristi C. Briggs, Jessica S. Lin, Lara Chaaban, Alyssa Parian, Mark Lazarev, Florin Selaru, Franck Housseau, Kellie N. Smith, Joanna M. P. Melia

## Abstract

**Background:** Ulcerative Colitis (UC) is characterized by chronic, relapsing and remitting inflammation in the colon and rectum. Pathogenic T cell activity is thought to play a major role in this process. T cell effector function is determined by the T cell receptor (TCR) and the antigen it recognizes. Examining the TCR repertoire can provide key insights into the adaptive immune response.

**Objective:** To characterize the longitudinal TCR repertoire of patients with UC across disease activity to determine if recurrent antigen(s) are responsible for active inflammation.

**Design:** Bulk TCR Vβ sequencing was done on colon tissue of 20 patients with UC across multiple time points of disease. Corresponding clinical metadata was also obtained over the same time period for each patient to map their clinical disease course. The top ten most highly abundant clones from each time point were longitudinally tracked and correlated with disease phenotype.

**Results:** Seventy-five percent of patients did not have overlapping abundant TCR clones across multiple time points of disease. The remaining 25% of patients had one to five TCR clones present in high abundance in their tissue during every time point analyzed.

**Conclusion:** These results demonstrate that most patients with UC do not share a similar TCR repertoire over time, indicating that times of inflammation are associated with unique antigen exposures. A smaller group of patients have persistent, private TCR clones with high abundance, 60% of whom had more unremitting, active disease.

## INTRODUCTION

Ulcerative colitis (UC) is one of two major forms of inflammatory bowel disease (IBD). There is a growing prevalence of the disease worldwide. UC affects about 1 in 250 people in North America, leading to estimates of 1.5 million people living with UC [1]. The typical cadence of disease in UC is notable for a waxing and waning course where there are times of active inflammation (flare) in the colon and rectum and times of low-level or absent inflammation (remission), but the amplitude of change between these time points is variable, and some patients can have sustained, unremitting disease [2, 3].

The pathogenesis of UC remains unknown, but it is hypothesized to be predominantly driven by pathologic T cell responses that lead to intestinal damage [3]. In murine models of colitis, T cell-mediated injury only occurs in the presence of a gut microbiota, implicating that T cells are activated in response to the resident microbiota and cause mucosal impairment [4]. T cell function is centered on the T cell receptor (TCR), which is a cell surface receptor comprised of an alpha and beta chain that recognizes peptides bound to the major histocompatibility complex (MHC). The specificity of this interaction relies on the amino acid sequence of the TCR which is encoded through VDJ recombination that results in a diverse repertoire in humans that can recognize virtually any antigens [5]. The antigens responsible for T cell responses in UC remain relatively unknown. Circulating and intestinal CD4+ T cells have been found to be reactive to the intestinal microbiome in healthy controls and patients with IBD [6].

An individual T cell responds to a specific peptide (or antigen) so it is well-established that the antigenic experience of any given T cell can be determined by examining the TCR repertoire. In response to a cognate antigen, proliferating T cells are characterized by clonotypic expansion and a resulting oligoclonal TCR repertoire comprised of identical or highly related TCR Vα and Vβ genes are generated. This type of clonal expansion can be identified by sequencing the TCR repertoire. The estimated potential TCR diversity in humans is up to 10^20^ with a total number of T cells estimated to be about 10^11^, indicating the ability for T cells to cross-react with multiple peptide antigens [5]. Currently, several assays exist to examine the TCR repertoire through sequencing of the TCR gene and can give accurate information about the relative frequency of different TCRs in blood or tissue [5, 7]. Multiplex PCR from genomic DNA is the most frequently used commercial assay available and is considered the gold standard in the field. An advantage of this method is that it can be done from formalin fixed paraffin embedded (FFPE) samples. This type of analysis has been frequently done in the field of immuno-oncology to identify T cells in the tumor microenvironment, leading to the ability to track tumor evolution, predict response to therapy, and develop novel T cell-specific therapies [7].

The application of TCR sequencing has spread to other fields including autoimmunity and inflammatory disorders. Disease-associated TCRs have been found in Celiac disease, type 1 diabetes, lupus, and ankylosing spondylitis and can act as biomarkers for disease and even therapeutic targets [8–10]. Memory T cell clones are known to play a role as self-antigens in multiple autoimmune disorders. Recent work in ankylosing spondylitis showed specific elimination of a particular TCR sequence known as, TRBV9, in a patient with treatment refractory disease led to durable clinical remission [7, 11]. Emerging literature on targeting of specific TCR-MHC complexes may lead to a novel personalized therapy approach in treating immune mediated diseases, and therefore, should be further considered in IBD.

Although UC is chronic and episodic in nature, many translational human studies in UC have a cross-sectional design using tissue obtained at a single time point with samples often compared to healthy (normal) or un-inflamed tissue. These studies cannot address the central question, how similar or different is the T cell response in an individual patient over time? To address this research gap, we performed TCR Vβ sequencing for a real-world cohort of patients with UC across multiple time points. We found the longitudinal TCR repertoire remains private in individual patients, meaning the highest frequency T cell clones associated with each time point is essentially distinct, suggesting limited shared antigenic exposure over time. In contrast, in a small subset of patients, persistent TCR clones in high abundance were present over multiple time points. Patients in this subgroup tended to have more medically refractory disease resulting in active disease in many time points analyzed.

## RESULTS

We performed TCR Vβ sequencing on a cohort of 20 patients with UC from sigmoid colon biopsy tissue. In total, nearly 60 time points were analyzed. Patient characteristics are given in Table 1. Summary clone statistics including number of clones, maximum frequency, and clonality for each patient and corresponding time points analyzed with TCR sequencing are found in Table 2.

**Table 1:**
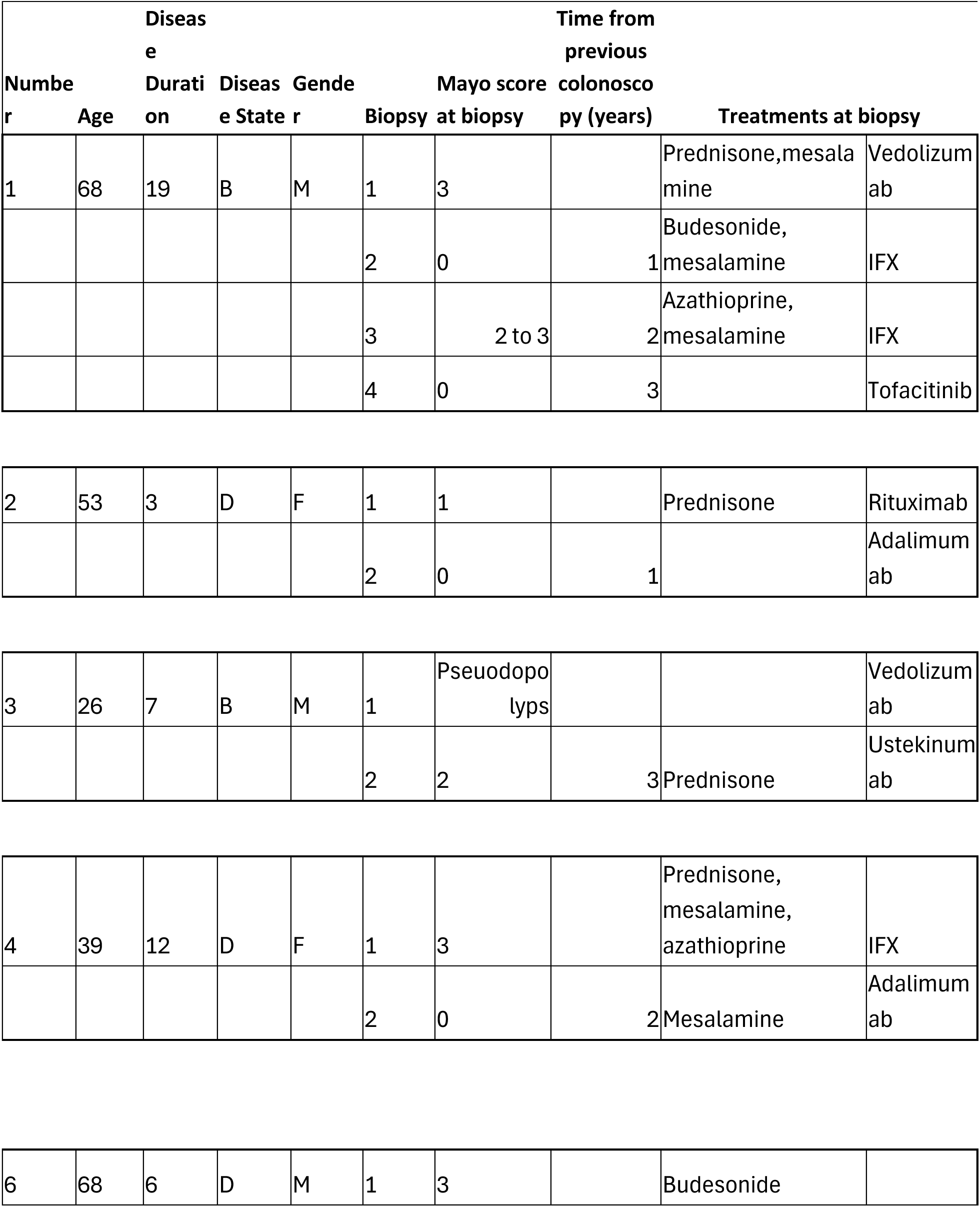
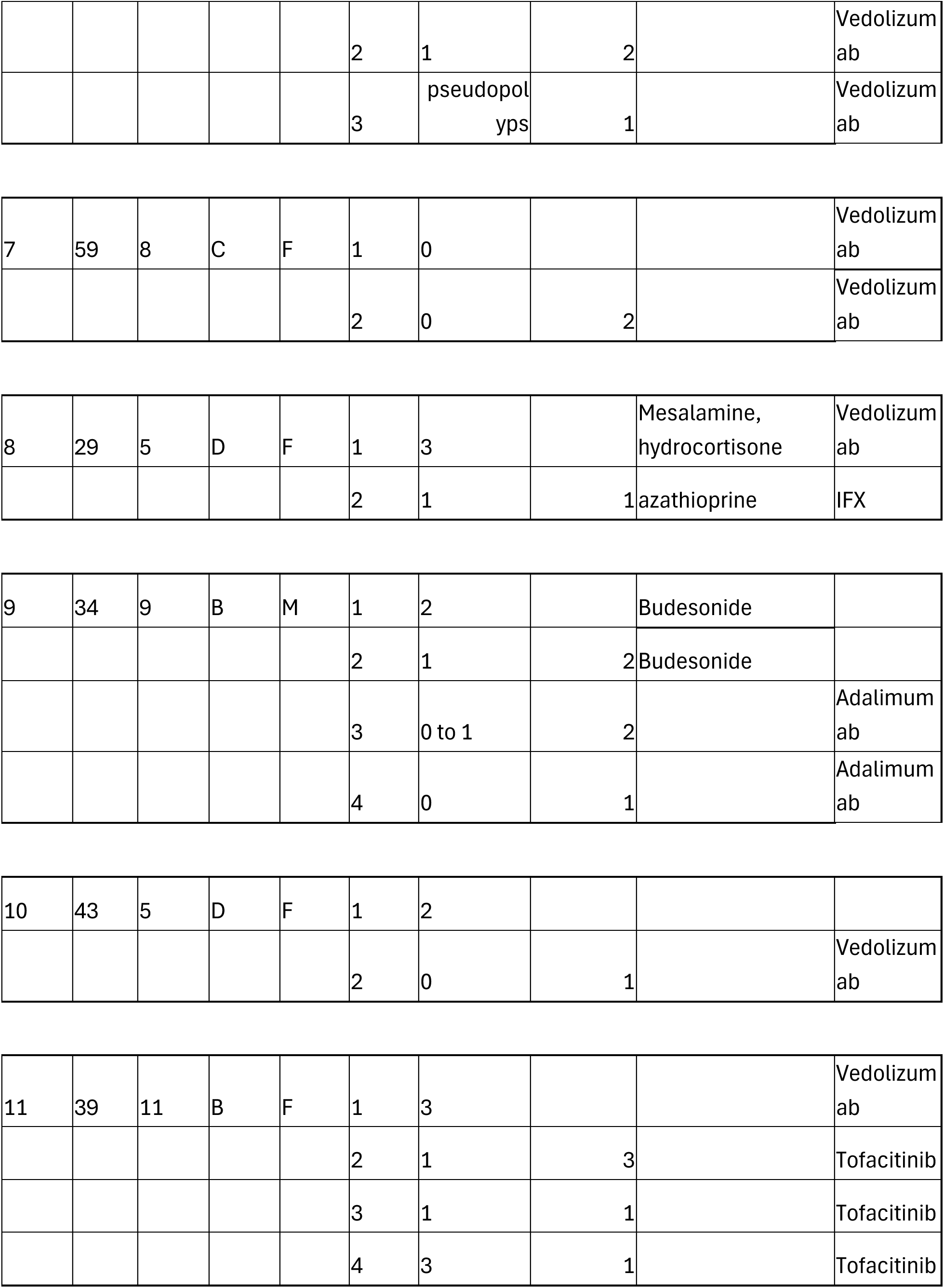
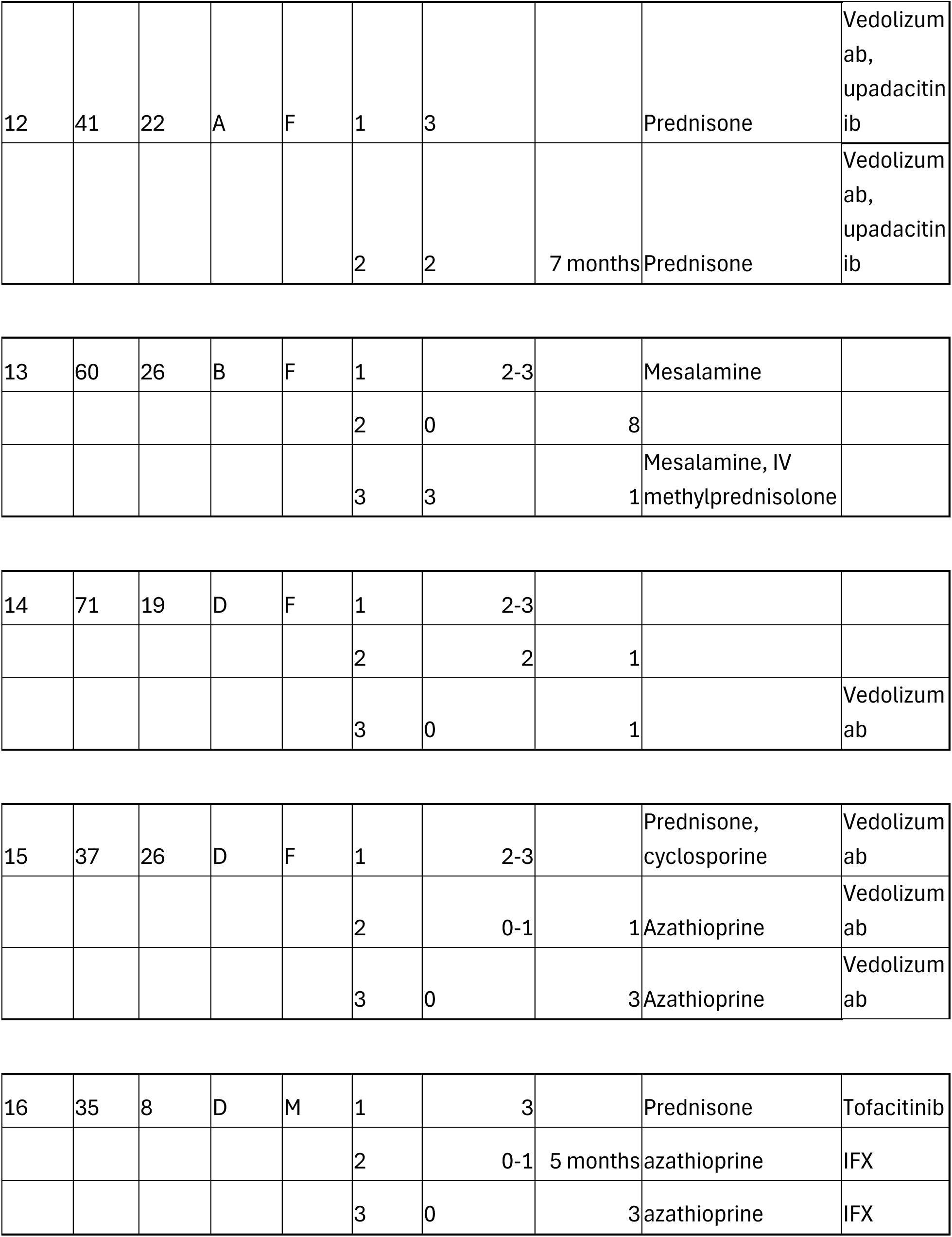
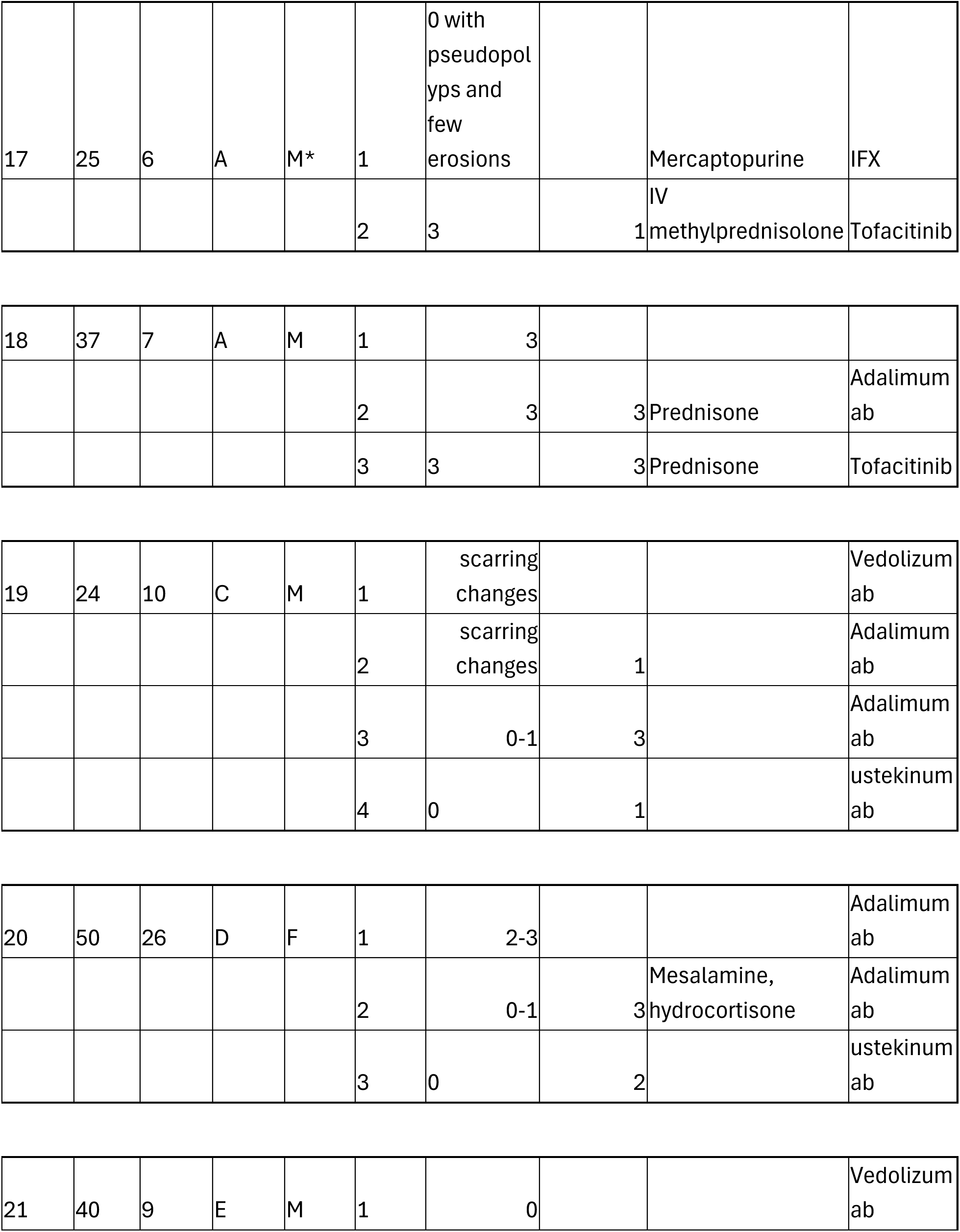
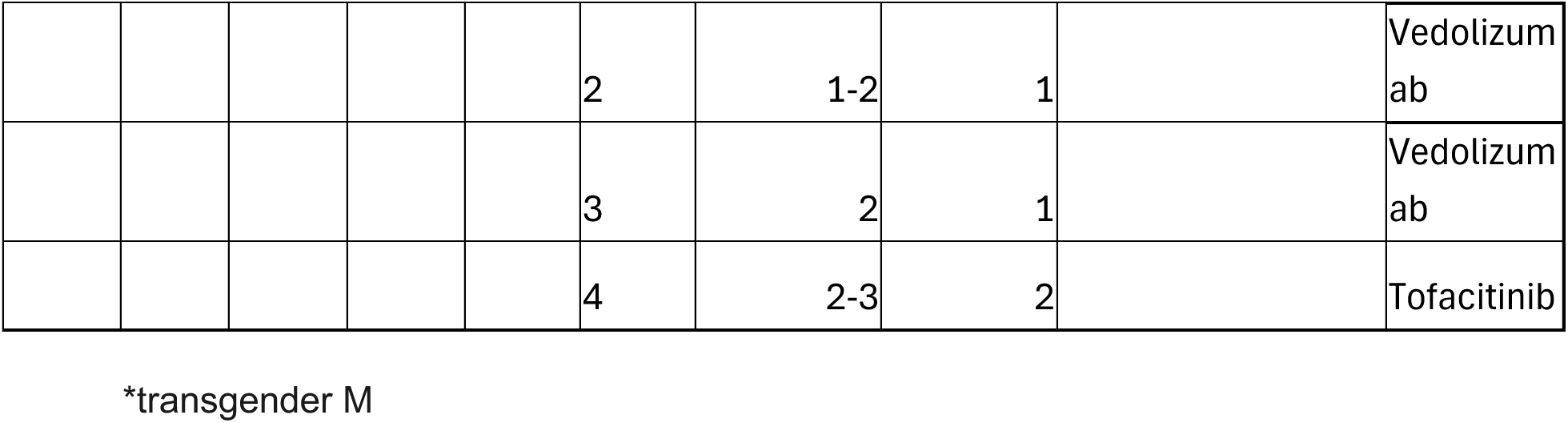
Patient characteristics.

**Table 2:** Clone summary statistics.

| **Patient Number** | **Time Point** | **Total Productive Reads** | **Maximum Productive Frequency** | **Total Clonotypes** | **Clonality** |
| --- | --- | --- | --- | --- | --- |
| 1 | 1 | 150691 | 0.0268 | 453 | 0.1969 |
| 1 | 2 | 142083 | 0.2435 | 164 | 0.3203 |
| 1 | 3 | 81030 | 0.0527 | 319 | 0.2446 |
| 1 | 4 | 143066 | 0.0895 | 132 | 0.2802 |
| 2 | 1 | 96712 | 0.1271 | 199 | 0.3940 |
| 2 | 2 | 160357 | 0.1115 | 299 | 0.2812 |
| 3 | 1 | 50325 | 0.0729 | 372 | 0.2713 |
| 3 | 2 | 166491 | 0.0576 | 1444 | 0.3956 |
| 4 | 1 | 196480 | 0.0421 | 537 | 0.2649 |
| 4 | 2 | 4519 | 0.0285 | 398 | 0.1576 |
| 5 | 1 | 31617 | 0.1735 | 167 | 0.4084 |
| 6 | 1 | 193880 | 0.0246 | 1070 | 0.2099 |
| 6 | 2 | 30724 | 0.0926 | 308 | 0.3233 |
| 6 | 3 | 117610 | 0.1531 | 352 | 0.3295 |
| 7 | 1 | 209959 | 0.0427 | 375 | 0.2053 |
| 7 | 2 | 196493 | 0.0759 | 316 | 0.3060 |
| 8 | 1 | 311913 | 0.0326 | 510 | 0.2087 |
| 8 | 2 | 340539 | 0.0442 | 4017 | 0.2097 |
| 9 | 1 | 411584 | 0.0198 | 1293 | 0.1849 |
| 9 | 2 | 90433 | 0.0739 | 155 | 0.2301 |
| 9 | 3 | 142848 | 0.0864 | 748 | 0.3041 |
| 9 | 4 | 283559 | 0.0127 | 2330 | 0.1524 |
| 10 | 1 | 90090 | 0.0299 | 1299 | 0.2420 |
| 10 | 2 | 70142 | 0.1221 | 351 | 0.3099 |
| 11 | 1 | 123744 | 0.0796 | 145 | 0.2171 |
| 11 | 2 | 273105 | 0.0130 | 588 | 0.1098 |
| 11 | 3 | 430527 | 0.0070 | 1938 | 0.1056 |
| 11 | 4 | 263154 | 0.0322 | 246 | 0.1409 |
| 12 | 1 | 783298 | 0.0402 | 4226 | 0.1556 |
| 12 | 2 | 487532 | 0.0265 | 3152 | 0.1367 |
| 13 | 1 | 20594 | 0.0716 | 396 | 0.2462 |
| 13 | 2 | 36699 | 0.0528 | 304 | 0.2255 |
| 13 | 3 | 240626 | 0.0220 | 1926 | 0.1115 |
| 14 | 1 | 22668 | 0.0444 | 275 | 0.2548 |
| 14 | 2 | 181677 | 0.0365 | 3325 | 0.1723 |
| 14 | 3 | 343099 | 0.0324 | 2083 | 0.1287 |
| 15 | 1 | 101014 | 0.0228 | 431 | 0.1979 |
| 15 | 2 | 409645 | 0.0081 | 1466 | 0.1241 |
| 15 | 3 | 493599 | 0.0456 | 1198 | 0.1265 |
| 16 | 1 | 279038 | 0.0494 | 305 | 0.1846 |
| 16 | 2 | 481724 | 0.0268 | 916 | 0.1390 |
| 16 | 3 | 252092 | 0.0598 | 253 | 0.2515 |
| 17 | 1 | 336002 | 0.0108 | 1922 | 0.1048 |
| 17 | 2 | 274710 | 0.0070 | 5958 | 0.0859 |
| 18 | 1 | 93592 | 0.0213 | 892 | 0.1258 |
| 18 | 2 | 538500 | 0.0168 | 5954 | 0.1087 |
| 18 | 3 | 827566 | 0.0197 | 4523 | 0.1023 |
| 19 | 1 | 42015 | 0.0646 | 358 | 0.2210 |
| 19 | 2 | 79771 | 0.0344 | 521 | 0.1910 |
| 19 | 3 | 308219 | 0.0051 | 2197 | 0.0984 |
| 19 | 4 | 410897 | 0.0107 | 4565 | 0.0849 |
| 20 | 1 | 276077 | 0.0092 | 1130 | 0.1234 |
| 20 | 2 | 80949 | 0.0167 | 772 | 0.1266 |
| 20 | 3 | 420233 | 0.0164 | 2889 | 0.0982 |
| 21 | 1 | 60953 | 0.0390 | 249 | 0.1787 |
| 21 | 2 | 100372 | 0.0362 | 518 | 0.1508 |
| 21 | 3 | 171447 | 0.0202 | 631 | 0.1473 |
| 21 | 4 | 100720 | 0.0123 | 822 | 0.1143 |

Based on histopathology and endoscopic appearance of disease (Mayo score) each time point was classified as either flare or remission. Clinical metadata including medication exposure, C-reactive protein (CRP), fecal calprotectin, and Clostridioides difficile or cytomegalovirus infection was also gathered from the electronic medical record at the time of the colonoscopy as well as intervening time points, when available (Table 3). As stated previously, the course of disease activity can be heterogenous in patients. As such, we defined distinct clinical phenotypes of disease activity in our cohort by evaluating all the clinical metadata available across time points, inspired by Solberg, et al [12]. Patients were divided into a total of five phenotypes labeled A through E (Figure 1A).

**Figure 1.**
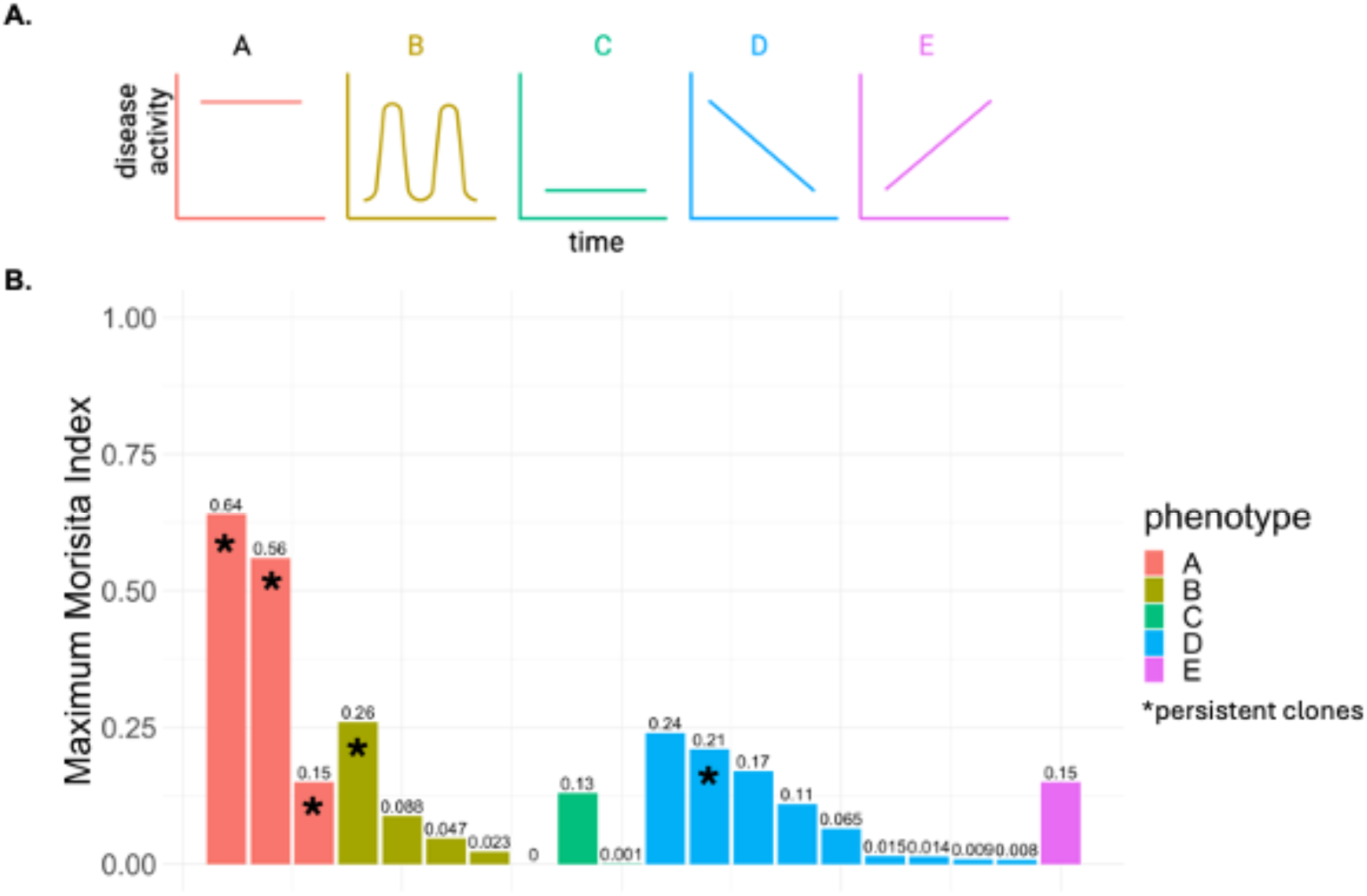
**A)** Disease phenotypes. Created with BioRender.com. **B)** average c reactive protein (CRP) and fecal calprotectin for all patients within each phenotype over the course of time examined by TCR sequencing. **C)** The maximum Morisita index (index range 0-1, with values closer to 1 indicating more overlap) of the TCR repertoire present amongst two time points of disease in each patient

**Table 3:** Clone overlap.

| **Patient** | **Number of Shared Clones in All Time Points** | **Number of time points** | **Time points with Highest Overlap** | **Maximum Morisita Index** |
| --- | --- | --- | --- | --- |
| 1 | 0 | 4 | 1F and 3F | 0.023 |
| 2 | 3 | 2 | F & R | 0.015 |
| 3 | 4 | 2 | F & F | 0 |
| 4 | 6 | 2 | F & R | 0.0081 |
| 5 |  |  |  |  |
| 6 | 8 | 3 | 1F and 3R | 0.17 |
| 7 | 7 | 2 | R & R | 0.00051 |
| 8 | 14 | 2 | F & R | 0.009 |
| 9 | 5 | 4 | F3 & R4 | 0.088 |
| 10 | 44 | 2 | F & R | 0.014 |
| 11 | 0 | 4 | 2R & 3R | 0.047 |
| 12 | 569 | 2 | F & F | 0.64 |
| 13 | 9 | 3 | 1F & 2R | 0.26 |
| 14 | 9 | 3 | 2F & 3R | 0.24 |
| 15 | 8 | 3 | 2R & 3R | 0.065 |
| 16 | 6 | 3 | 2F & 3R | 0.21 |
| 17 | 83 | 2 | F & F | 0.15 |
| 18 | 39 | 3 | 2F & 3F | 0.56 |
| 19 | 7 | 4 | 3F & 4R | 0.13 |
| 20 | 37 | 3 | 1F & R3 | 0.11 |
| 21 | 4 | 4 | 2F & 4F | 0.15 |

F = flare R = remission

Clonal T cell overlap across time points was first analyzed in an unbiased manner in all patients longitudinally. One method of quantifying TCR repertoire overlap is the Morisita index, where values closer to 1 indicate higher overlap. Within each patient, pair-wise comparisons were done between time points to calculate each Morisita value. The maximum Morisita index (or if only 2 time points were available, the only Morisita index) for each patient grouped based on disease phenotype is shown in Figure 1B. The total number of overlapping clones and specific time points corresponding to the maximum Morisita index is shown in Table 4.

The two highest Morisita index values are seen in patients with phenotype A, which is associated with high disease activity across time points. The highest overlap between any two samples had a Morisita index of 0.64 and a total of 569 overlapping T cell clones between two flares. Clinical data of this patient shows elevated fecal calprotectin and CRP values indicating high, sustained disease activity, which was also seen on endoscopy (Figure 2A). The average CRP and fecal calprotectin were 4925 µg/g and 13.78 mg/L, respectively, in this patient. Four of the top ten most abundant TCRs in time point one was also present in top ten abundance in time point two (Figure 2B & C, Supplementary Table 1). The second highest Morisita index of 0.56 was also in a patient with phenotype A who had 39 clones shared across all three time points. Four clones were also persistent in the top 10 most abundant T cell clones in all time points (data not shown). The third patient in phenotype A had 2 shared TCR clones in high abundance across flares. Therefore, all patients in phenotype A had persistent clones with variable time between colonoscopies, ranging from 7 to 12 months.

**Figure 2.**
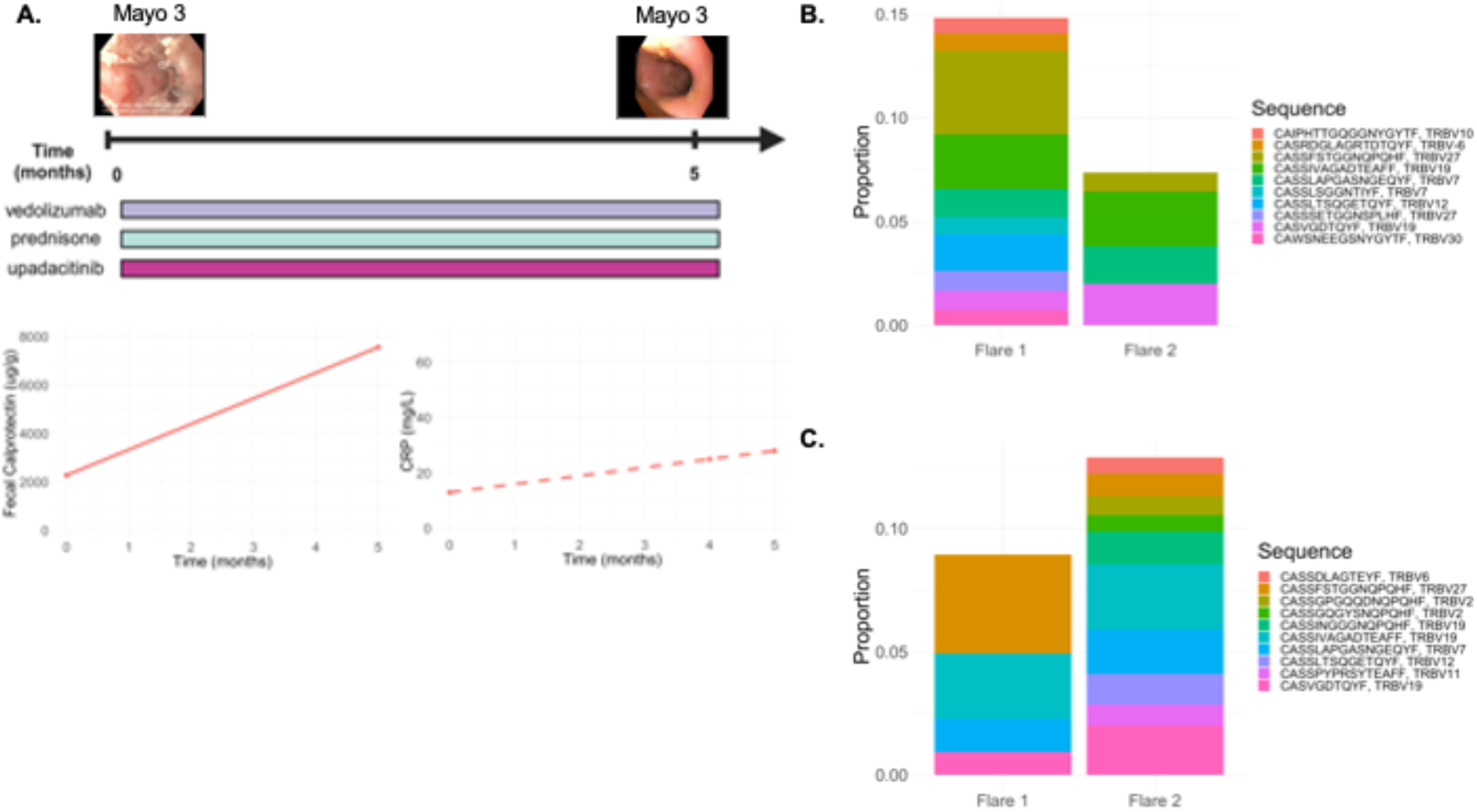
**A)** Serial colonoscopy and associated TCR sequencing time points with representative image of sigmoid colon and Mayo endoscopic severity score at location of image (top). Longitudinal clinical course of patient with medication administration (middle) and fecal calprotectin and CRP values (bottom). Time is presented as weeks following first colonoscopy/TCR sequencing analysis (time 0). Created with BioRender.com Top 10 most frequent clones at time point 1 (**B**) and time point 2 (**C**) and the associated frequency of these clones at the other time points. Only clones that are present in the top 10 most frequent clones are plotted for any time point.

The third highest Morisita index (0.26) was seen in a patient with a more relapsing and remitting form of the disease (Phenotype B). Two clones in the top 10 were present in high abundance across all four time points for this patient. However, the remaining four patients in phenotype B showed more heterogenous TCR repertoires. Figure 3A shows a patient with fluctuating fecal calprotectin and CRP. When the top ten most abundant T cell clones were tracked over time in this patient, no shared clones were present at other time points (Figure 3B and C, Supplementary Table 2). Two other patients in phenotype B also did not have any shared T cell clones across all time points analyzed (Table 4). Longitudinal TCR sequencing was done from tissue obtained from colonoscopies spanning 3-9 years in this group. Similarly, the two patients who started with low disease activity that remained in remission (phenotype C) also did not show T cell clones that persisted in high abundance over multiple time points (Figure 4, Supplementary Table 3).

**Figure 3.**
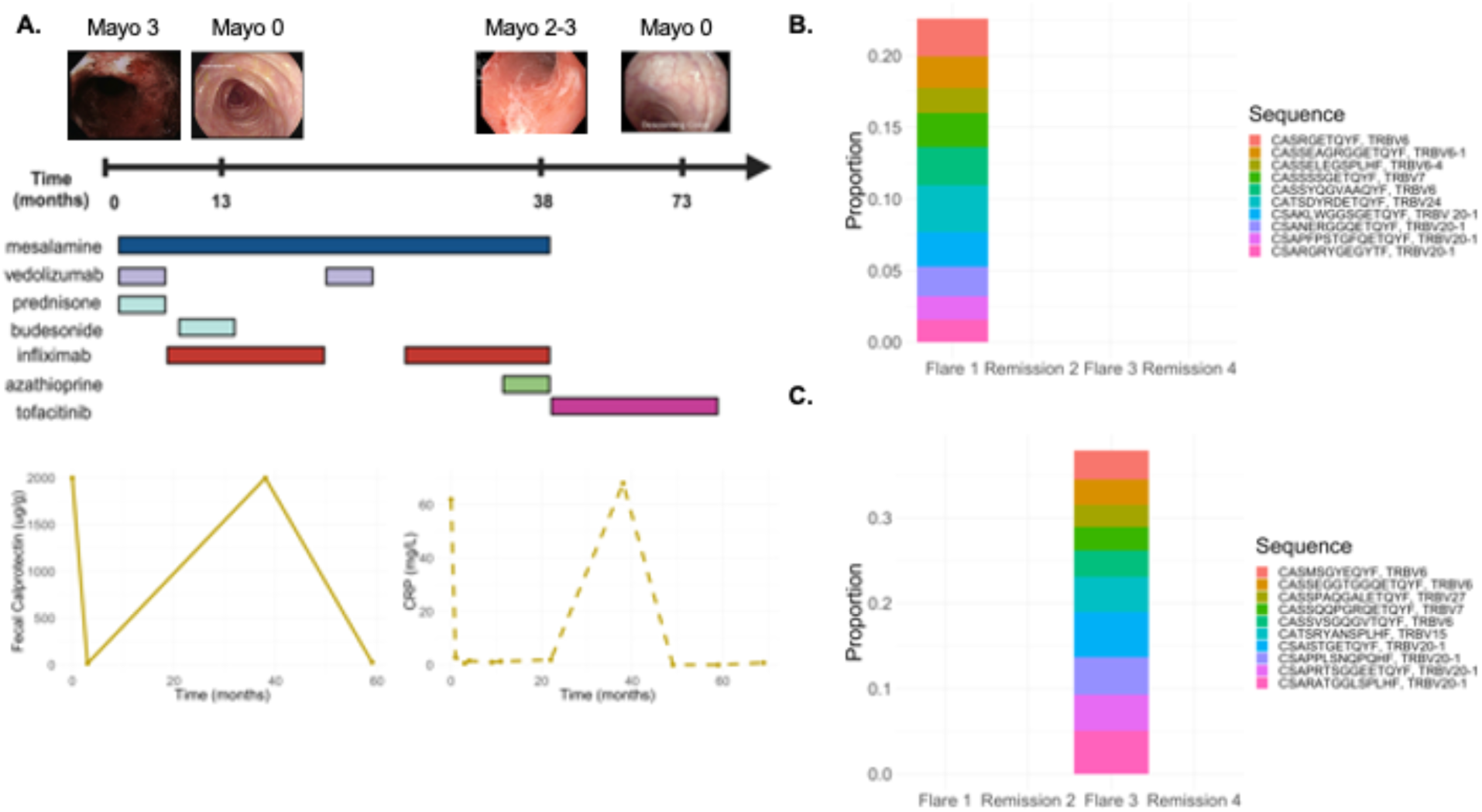
**A)** Serial colonoscopy and associated TCR sequencing time points with representative image of left colon and Mayo endoscopic severity score at location of image (top). Longitudinal clinical course of patient with medication administration (middle) and fecal calprotectin and CRP values (bottom). Time is presented as weeks following first colonoscopy/TCR sequencing analysis (time 0). Created with BioRender.com Top 10 most frequent clones at time point 1 (**B**) and time point 3 (**C**) and the associated frequency of these clones at the other time points. Only clones that are present in the top 10 most frequent clones are plotted for any time point.

**Figure 4.**
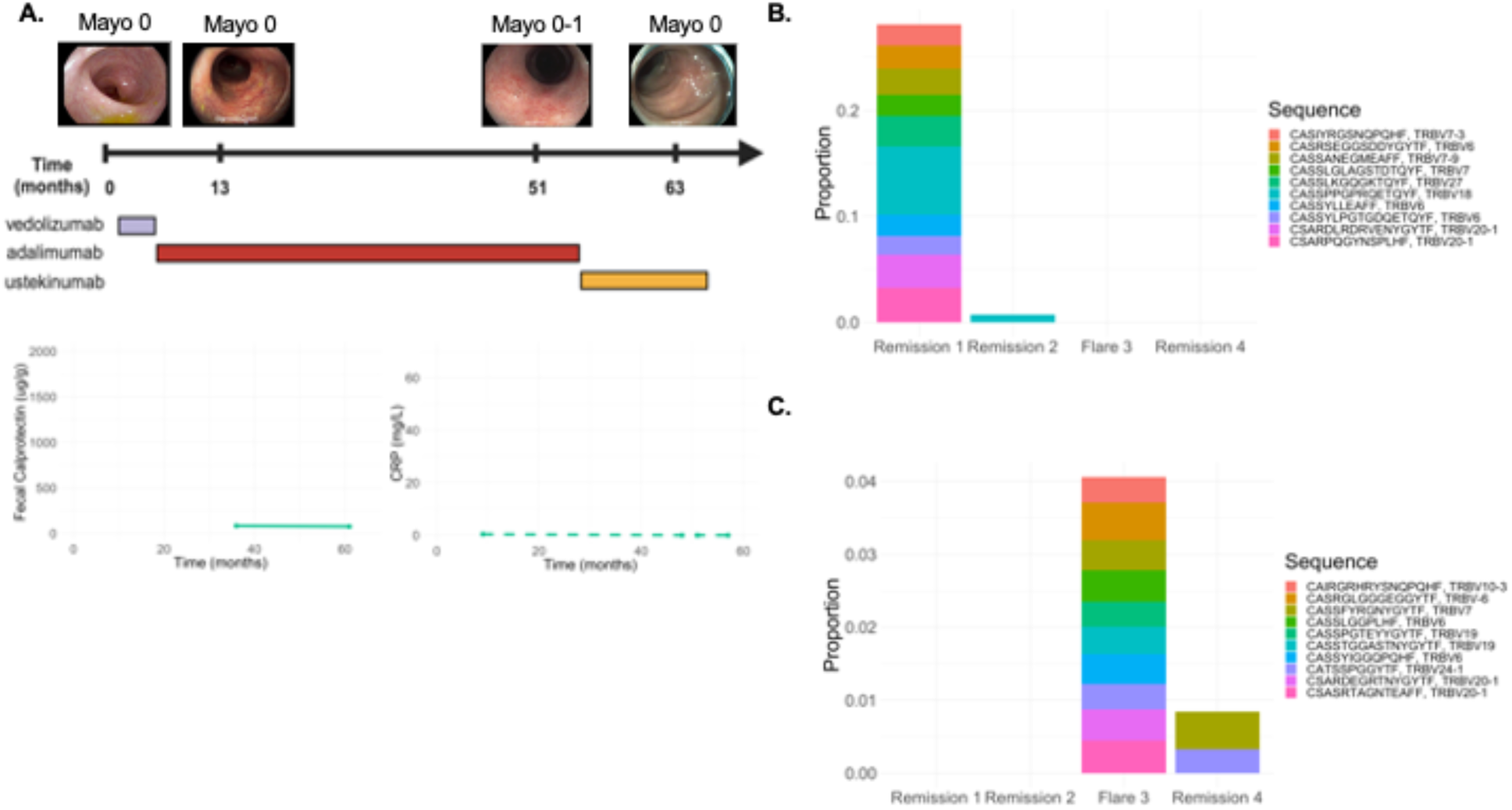
**A)** Serial colonoscopy and associated TCR sequencing time points with representative image of sigmoid colon and Mayo endoscopic severity score at location of image (top). Longitudinal clinical course of patient with medication administration (middle) and fecal calprotectin and CRP values (bottom). Time is presented as weeks following first colonoscopy/TCR sequencing analysis (time 0). Created with BioRender.com Top 10 most frequent clones at time point 1 (**B**) and time point 3 (**C**) and the associated frequency of these clones at the other time points. Only clones that are present in the top 10 most frequent clones are plotted for any time point.

Forty-five percent of the patient cohort were phenotype D, representing high disease activity followed by remission. The highest Morisita index in this group was 0.24 (Figure 1C), which was found between a flare and remission time point. At all 3 time points analyzed in this patient, there were no persistent high abundance clones. The patient with a Morisita index of 0.21 in phenotype C had one shared TCR clone in the top 10 over four time points. The remaining seven patients in phenotype D had distinct TCR repertoires with minimal overlap of clones across time points. Clinical data from a patient in this group shows high CRP and fecal calprotectin that then normalized (Figure 5A). One of the top ten clones found in time point one was also present in high abundance in time point two. Two clones from time point three were also present in the top ten of time point four (Figure 5B & C, Supplementary Table 4), but again, no clones were in the top ten of all time points. Time interval between colonoscopies in this group ranged from 5 months to 3 years.

**Figure 5.**
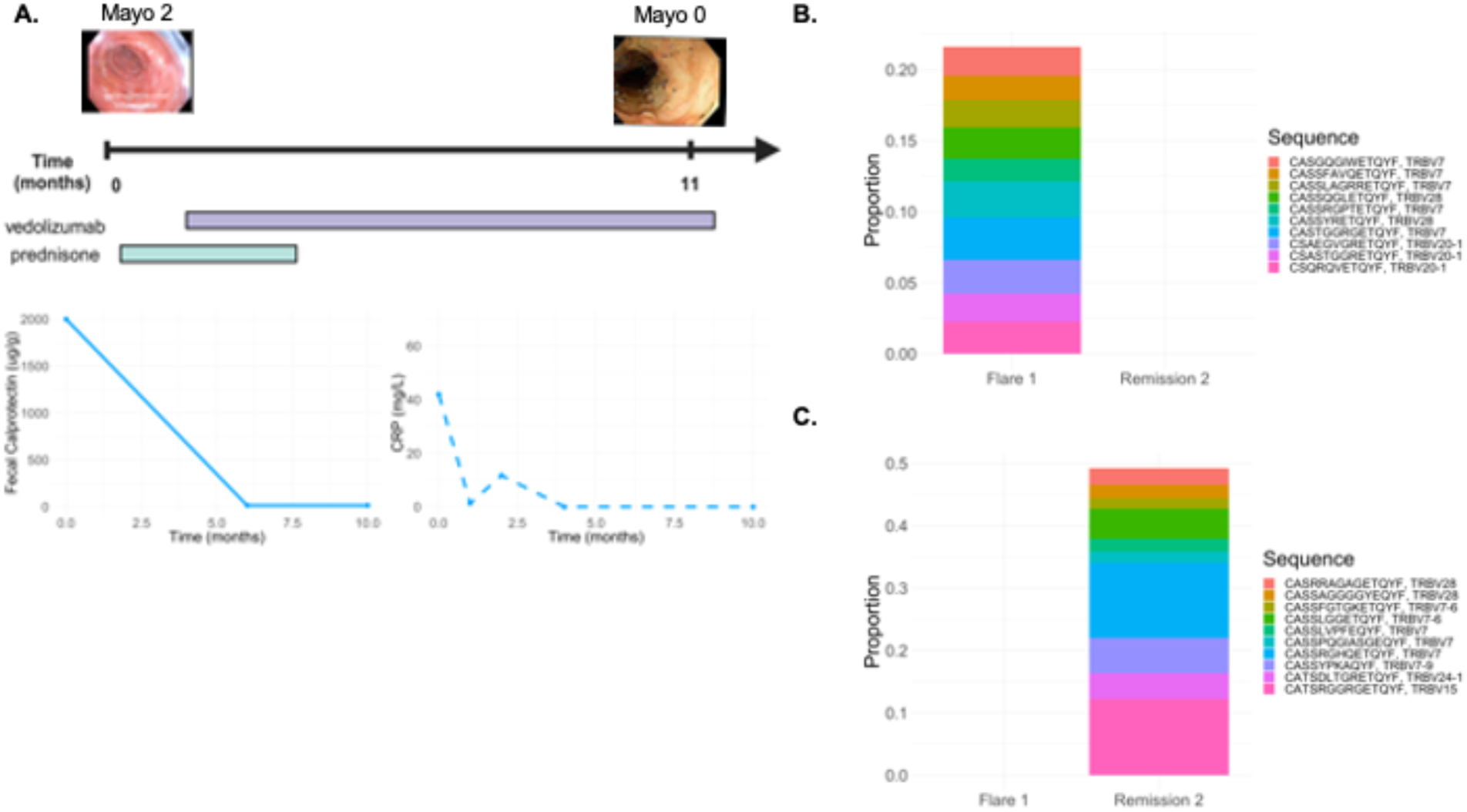
**A)** Serial colonoscopy and associated TCR sequencing time points with representative image of sigmoid colon and Mayo endoscopic severity score at location of image (top). Longitudinal clinical course of patient with medication administration (middle) and fecal calprotectin and CRP values (bottom). Time is presented as weeks following first colonoscopy/TCR sequencing analysis (time 0). Created with BioRender.com Top 10 most frequent clones at time point 1 (**B**) and time point 2 (**C**) and the associated frequency of these clones at the other time points. Only clones that are present in the top 10 most frequent clones are plotted for any time point.

Lastly, one patient had low disease activity and then flared (phenotype E). TCR repertoire analysis was similar to phenotypes B, C, and D where repertoires were distinct at each time point (with 1-2 years between each colonoscopy). The same clones were not present in high abundance across all time points, although, 1-2 clones were shared in high abundance between two of the three flare time points.

## DISCUSSION

In this study, we examine TCR repertoires to longitudinally immunophenotype patients with UC, a critical gap in the current literature. The use of TCR sequencing has been a well-established mechanism to survey the overall clonal T cell populations in tissue as well as the TCR repertoire present in a sample. These methods have been validated and used in cancer immunology and other diseases extensively [5, 7]. We used a real-world data set in this study with a cohort of patients on a variety of medications, including advanced therapies and steroids, with a spectrum of disease activity. The longitudinal TCR repertoire of patients over time was heterogenous in most patients. Therefore, the most abundant clones associated with each time point were distinct, thus indicating that a unique set of antigens are activating T cells at each flare. In a subset of patients (25%), highly abundant T cell clones were identified that were present across all time points. All patients with a medically-refractory, unremitting phenotype (A) had persistent clones. The significance of these findings for how we frame disease initiation and persistence in UC are two-fold. First, in patients with a relapsing and remitting disease activity phenotype, the TCR repertoire supports a model in which flares are immunologically distinct rather than a model whereby there is recurrent antigenic stimulation prompting the flare. Second, in patients with unremitting disease, there is evidence of shared antigenic exposure given shared, high abundance T cell clones, potentially representing an important therapeutic target.

Most studies applying TCR sequencing in IBD have focused on comparing TCR repertoires in patients with IBD to normal healthy controls or adjacent un-inflamed tissue to find public (present in more than 1 individual) TCRs in IBD patients. In these analyses, expanded CD4+ T cell clones have been found in the inflamed tissues of Crohn’s Disease (CD) patients with one study showing up to 58% expansion of a single clone [13–15]. Oligoclonal expansion of T cells have also been found in the normal intestine, but not to the same degree as disease [16, 17].

The overall TCR repertoire diversity has been shown to decrease in CD compared to healthy controls [18]. In CD, the preferential use of certain TCRs have been reported in a subset of patients, but the identity and abundance of the public TCRs is variable across manuscripts [19–21].

The concept of persistent T cell clones has been presented before in the literature in patients with UC. Chott, *et al*., 1996 showed the presence of TRBV3 to be preferentially abundant in patients with UC, specifically those who went on to colectomy [16]. A follow-up study (Probert, *et al*., 2001) then analyzed this clone during active disease and remission and saw the presence of this clone in both time points in patients, indicating that it was not correlated with disease activity, so the longitudinal significance of this clone was questioned [22]. We too found that 11 out of 20 patients in our cohort had TRBV3 T cell gene usage without correlation with disease state. In Chott, *et al*, the persistent clones were first identified in patients who had medically refractory disease requiring colectomy. Consistent with this observation, in our data set, three of the five patients who had persistent T cell clones were also the patients in need of colectomy (though one did not proceed due to personal preference). Additionally, expanded clones have remained in the peripheral blood of patents with CD for up to a year, suggesting there is a sustained response to a particular antigen during chronic inflammation [19]. Taken together, these data emphasize that we have an incomplete understanding of the clinical significance of T cell repertoires, but the association with more refractory disease found now and in the historical literature supports reconsideration of the role these clonal populations play in UC.

In individuals who have disease that is medically refractory, our data opens the door for the possibility of TCR sequencing revealing a private, persistent clone that could act as a therapeutic target or a biomarker of disease. These types of clones suggest that there may be discrete antigen(s) that are specific to a particular patient or at least a smaller cohort of patients. Multiple gut commensal organisms can signal to the same T cell, therefore, a common T cell clone could be persisting due to a group of organisms causing a pathologic immune response in a susceptible host that may not be adequately targeted with therapy [23]. A higher overlap of TCRs in the blood and intestine has been shown in IBD compared to healthy individuals so persistent TCRs may also appear in the peripheral blood in patients [24, 25]. Finally, recent data shows CD8 T cells are spread homogenously in the colon with high repertoire overlap between locations indicating sample location should not interfere with these results [25].

The technology to specifically target TCR clones has improved tremendously in the past couple decades, especially with the advent of CAR-T cell therapies. In addition, the concept of a pathologic T cell clone has gained more attention presently as specific targeting of TRBV9 resulted in clinical remission in a patient with medically refractory ankylosing spondylitis. Antibody-mediated depletion of TRV9 positive T cells was successfully administered to the patient with demonstrated safety and efficacy [11]. There is also existing evidence in the literature for the use of TCR clones as a predictive biomarker. Following therapy with an anti-TNF, responders to treatment showed larger changes in their TCR repertoire when compared to non-responders where there was a smaller number of overlapping clones in non-responders. Although the repertoire remained largely unchanged in both groups [15]. The difference seen between responders and non-responders was present in the most expanded clones of the repertoire, supporting the notion that the highest abundant clones are likely the most relevant. Another study found CD4+ NKG2D+ T cells were predictive of post-operative relapse in CD [26]. A TRBV10-1-TRBJ2-7 clonotype significantly correlated with a higher Rutgeerts score among patients with post-operative recurrence of CD [27]. Given these findings in CD, one could imagine targeting specific TRBV clones in UC that are found to be associated with or cause refractory inflammation could result in successful remission as seen in ankylosing spondylitis.

A number of limitations exist in this study, which include the inability to identify the specific type of T cell associated with a given clone, our focus on the high frequency clones, and limited patient sample size. CDR3 sequence amino acid sequences were queried on VDJdb and McPAS to determine the identity of TCR clones [28]. One of the persistent overlapping clones had sequence that matched an epitope against influenza A, the remaining did not have any matches. A total of 14 abundant clones from all patients could be identified and were found to be against cytomegalovirus, insulin-like growth factor, influenza A, and Epstein-Barr virus. Viral reactive clonal T cells are known to be common in IBD [29]. Literature also shows the expansion of CD8 T cell clones in UC, thus, we could hypothesize that the dominant clones are likely CD8 T cells. Identification of these T cell clones is essential to answer the fundamental question that remains in this disease: What are the pathologic T cells in IBD that cause inflammation and how do they change over time? Even though our data indicates different antigenic signaling may be responsible for each flare in patients, our findings do not necessarily rule out the possibility of a “super antigen” that could be triggering multiple T cell responses or a group of antigens triggering an inflammatory response, which has been seen in autoimmunity. Future work is required to also focus on the lower frequency clones to determine their functional role. Expansion of our results is required in a larger cohort of patients, especially those who have persistently active disease compared to those who are maintained in remission, to determine the clinical applicability of our initial observations. Lastly, better understanding of the antigen that is being recognized by the expanded TCRs is essential in understanding the biological basis of this disease. Application of single cell analyses over a longitudinal cohort would allow for sequencing of both alpha and beta chains, identification of the associated T cell subtypes and cell states, and predictive modeling of MHC-dependent TCR-antigen interactions [30, 31].

## METHODS

### Patients and Clinical Data

FFPE blocks from patients were obtained as part of an Institutional Review Board (IRB) approved protocol for TCR sequencing (IRB 00336979). Blocks were obtained from the Johns Hopkins Hospital pathology archives. Patients who had archived colonoscopy biopsy specimen from time points of active disease and remission based on histopathology as documented by pathologists were selected for the study. A total of 23 patients were identified for this study and a total of 21 were used in the final analysis; two patients were dropped due to lack of sufficient reads during sequencing and lack of availability of paraffin block from the colonoscopy. In addition, some time points within patients were also dropped due to insufficient sequencing reads or block availability. Due to this patient 5 was not included in the analysis as only one time point remained. Ultimately, 20 patients with 2 to 4 time points were included in the study (Table 1). Definition of flare and remission were made based on both histologic and endoscopic data at time of colonoscopy. A flare was defined as active disease on pathology and Mayo score of 2 or 3 on endoscopy. Mayo score of 0 with pathology of inactive disease or normal colonic mucosa was considered to be remission. In rare circumstances where there was discrepancy between pathology and endoscopic findings, the cases were re-reviewed with a pathologist and classified based on their recommendation. Clinical data was also extracted from the electronic medical record including medication administration, endoscopy findings, pathology results, CRP, fecal calprotectin, medications, and infections including *Clostridioides difficile* or cytomegalovirus. All clinical data was collected based on available information in the medical record between the first time point and last time point of TCR tissue analysis for each patient. Patients were all subjectively assigned to a phenotype (A-E) based on disease activity as defined by the criteria above. Documentation of consent was waived by the IRB for this study, but verbal consent was obtained from each patient. Patient privacy was protected during data acquisition and storage as only members on the study team were given personalized level data and remaining data was anonymized. This manuscript does not contain any protected health information for any patient.

### TCR Sequencing & Analysis

DNA extraction from FFPE-preserved colon biopsy specimens was performed using the DNeasy Blood and Tissue Kit (Qiagen). The TCR-B locus was amplified and sequenced by the Johns Hopkins Sidney Kimmel Comprehensive Cancer Center FEST and TCR Immunogenomics Core Facility using the Ampliseq for Illumina TCR-short read kit. TCR sequencing and quality control was completed as previously described [32]. Briefly, non-productive TCR CDR3 sequences (premature stop or frameshift), sequences with amino acid length less than 7, and sequences not starting with ‘C’ or ending with ‘F/W’ were excluded from the final analyses. Specimens with at least 1000 reads were included in the final analysis. The degree of clonality for each specimen was assessed by the productive clonality matrix, which is defined as 1-Pielou’s evenness. Values near one represent samples with one or a few predominant clones (monoclonal or oligoclonal samples), whereas values near 0 represent a polyclonal population. Immunarch R package was used to analyze clonotype number, proportion, diversity, overlap, and tracking clones [33]. The degree of T cell clone overlap at the species level was evaluated using the Morisita overlap index. In values near one, the species occur in the same proportion in both samples, whereas values near 0 implies the two samples do not overlap in terms of species. Immunarch was also used to annotate clonotypes using the McPAS and VDJdb databases [33, 28].

### Data Availability

Pre-processed and de-identified TCR sequencing data will be deposited to a public repository with unrestricted access upon publication.

## PATIENT & PUBLIC INVOLVEMENT

Patients were involved in this study as human tissue from patients were the primary sample source in this project. Although tissue for this study was collective retrospectively so no additional burden was present on the patient population, each patient was individually contacted to get their verbal consent to participate (consent documentation was waived by the IRB). In addition, clinical data from the electronic health record of patients were collected in this study.

Authors of this study are clinical gastroenterologists who care for patients with IBD, including those included in this cohort. They are highly attuned to the clinical gaps in the field and formulate research questions based on their ongoing interactions with patients.

## ACKNOWLEDGEMENTS

We would like to acknowledge the TCR sequencing and FEST core led by Dr. Kellie Smith at the Johns Hopkins Bloomberg∼Kimmel Institute for Cancer Immunotherapy. We would like to acknowledge the Institutional Review Board at Johns Hopkins and thank all the patients who willingly participated in this study. We would also like to acknowledge Dr. Drew Pardoll and Dr. Cynthia Sears for their intellectual input in the study design and data analysis of this project. Funding for this study was provided by investigator-initiated research funded provided by Pfizer (J.M.), Bloomberg∼Kimmel Institute for Cancer Immunotherapy (K.N.S.), National Institutes of Health grant R37CA251447 (K.N.S.) and DK114478 (J.M.).

## ETHICS STATEMENT

This study involves the human participants and was approved by the an Institutional Review Board (IRB 00336979). This study does not involve animal subjects.

## CONFLICT OF INTEREST

There are no competing interest for any authors.

---

**Supplementary Table 1:**
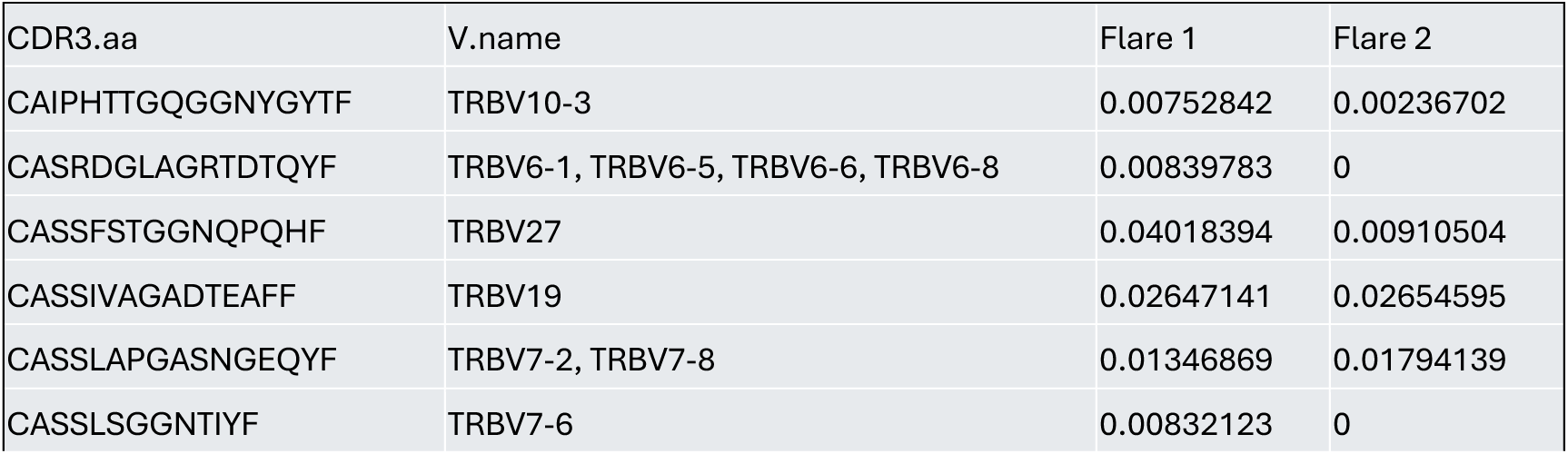
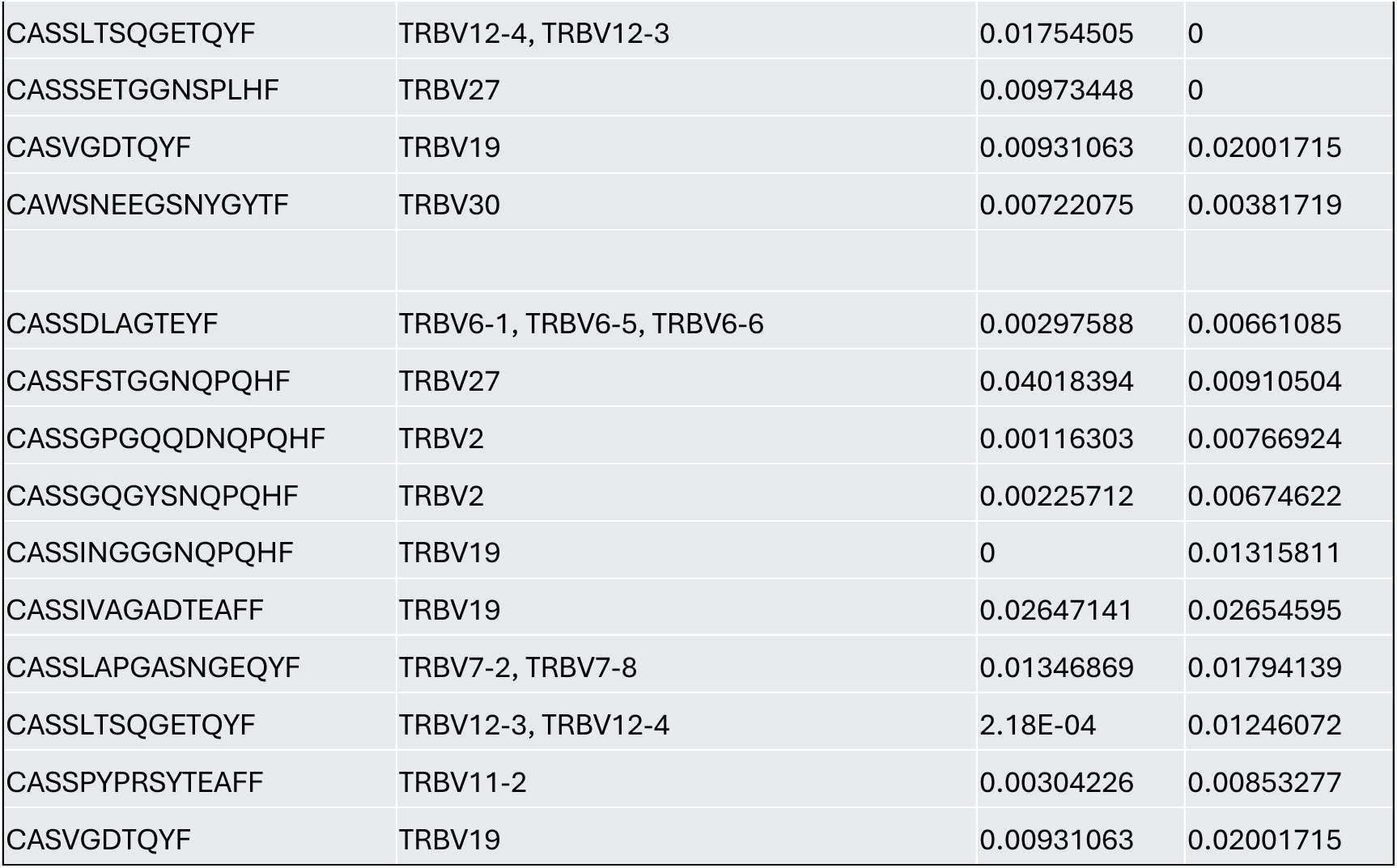
Frequency of top 10 clones in each time point.

**Supplementary Table 2:**
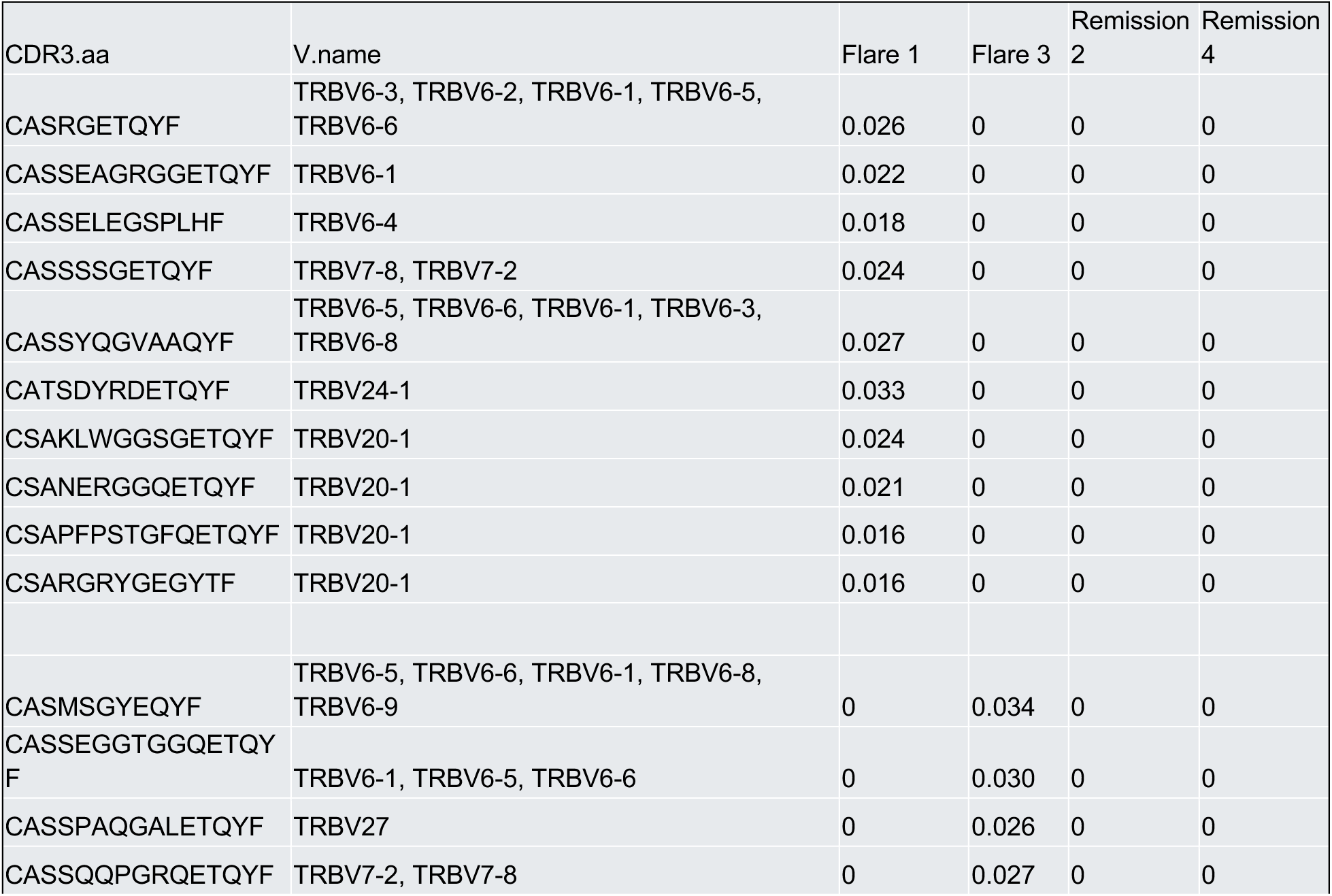
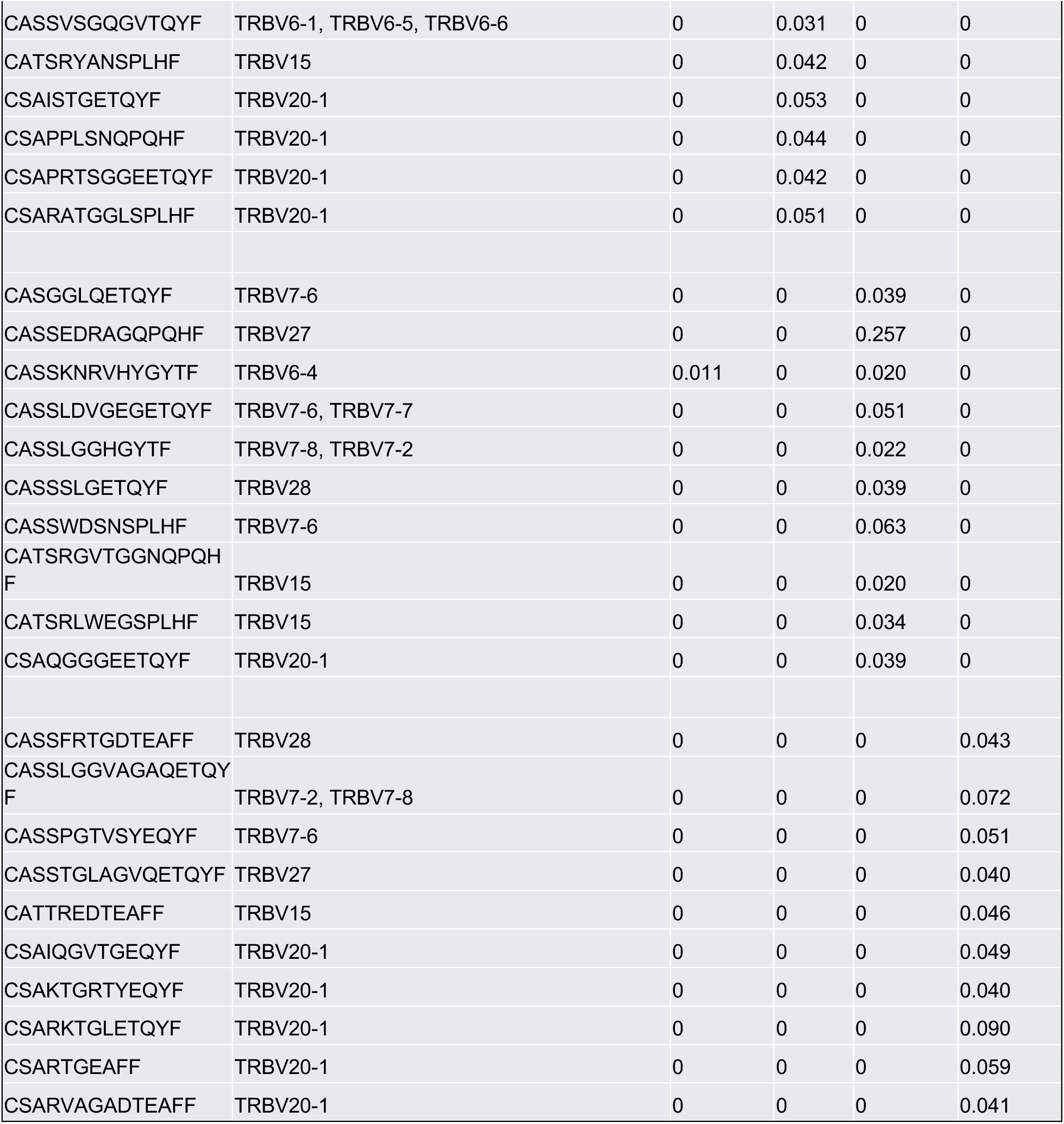
Frequency of top 10 clones in each time point.

**Supplementary Table 3:**
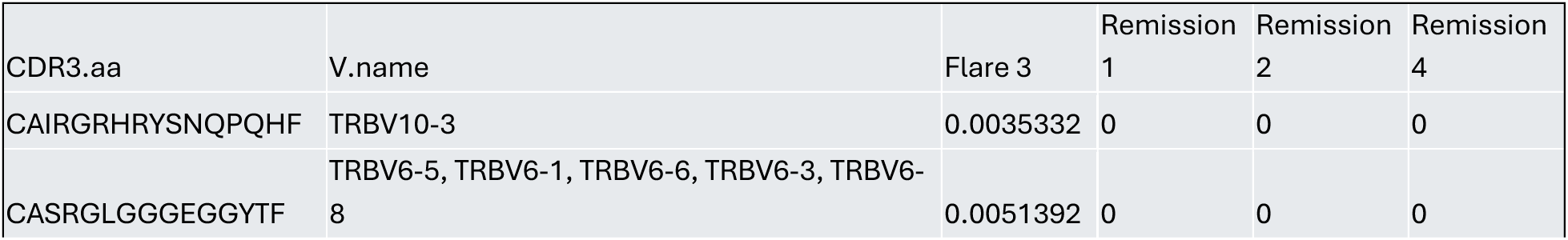
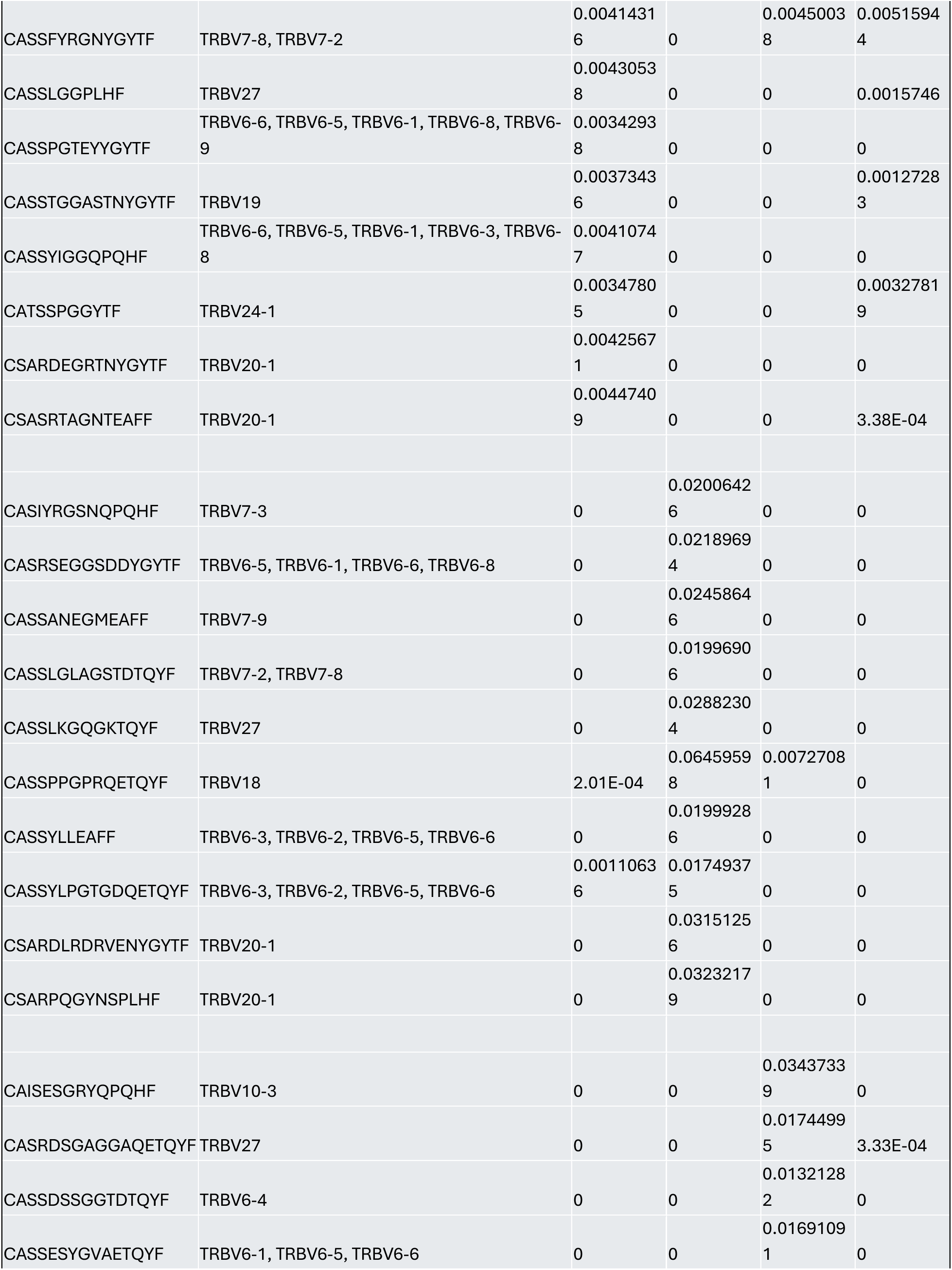
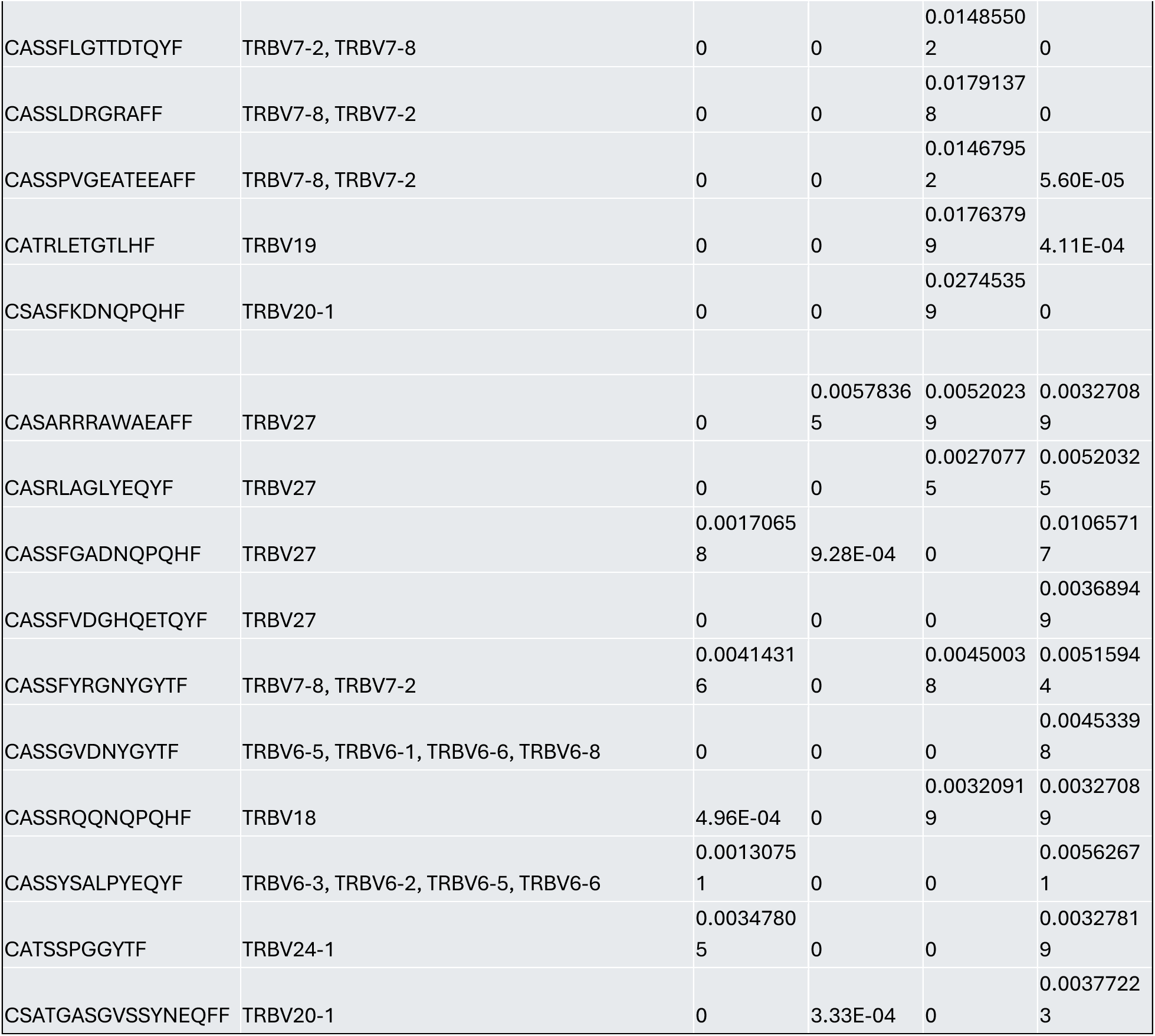
Frequency of top 10 clones in each time point.

**Supplementary Table 4:** Frequency of top 10 clones in each time point.

| CDR3.aa | V.name | Flare 1 | Remission 2 |
| --- | --- | --- | --- |
| CASGQGIWETQYF | TRBV7-2, TRBV7-8 | 0.02051282 | 0 |
| CASSFAVQETQYF | TRBV7-2, TRBV7-8 | 0.01691642 | 0 |
| CASSLAGRRETQYF | TRBV7-2, TRBV7-8 | 0.01926962 | 0 |
| CASSQGLETQYF | TRBV28 | 0.02224442 | 0 |
| CASSRGPTETQYF | TRBV7-2, TRBV7-8 | 0.01585082 | 0 |
| CASSYRETQYF | TRBV28 | 0.02533023 | 0 |
| CASTGGRGETQYF | TRBV7-2, TRBV7-8 | 0.02991453 | 0 |
| CSAEGVGRETQYF | TRBV20-1 | 0.02369852 | 0 |
| CSASTGGRETQYF | TRBV20-1 | 0.01989122 | 0 |
| CSQRQVETQYF | TRBV20-1 | 0.02251082 | 0 |
| CASRRAGAGETQYF | TRBV28 | 0 | 0.02677426 |
| CASSAGGGGYEQYF | TRBV28 | 0 | 0.02175587 |
| CASSFGTGKETQYF | TRBV7-6 | 0 | 0.01633829 |
| CASSLGGETQYF | TRBV7-6 | 0 | 0.04804539 |
| CASSLVPFEQYF | TRBV7-2, TRBV7-8 | 0.004662 | 0.02091472 |
| CASSPQGIASGEQYF | TRBV7-2, TRBV7-8 | 0 | 0.01783525 |
| CASSRGHQETQYF | TRBV7-6, TRBV7-7 | 0 | 0.12044139 |
| CASSYPKAQYF | TRBV7-9 | 0 | 0.05674204 |
| CATSDLTGRETQYF | TRBV24-1 | 0 | 0.04125916 |
| CATSRGGRGETQYF | TRBV15 | 0 | 0.12209518 |

## Notes

### Competing Interest Statement

The authors have declared no competing interest.

